# A versatile system to introduce clusters of genomic double-strand breaks in large cell populations

**DOI:** 10.1101/2020.04.27.064105

**Authors:** Thorsten Kolb, Umar Khalid, Milena Simović, Manasi Ratnaparkhe, John Wong, Anna Jauch, Peter Schmezer, Agata Rode, Shulamit Sebban, Daniel Haag, Michaela Hergt, Frauke Devens, Yosef Buganim, Marc Zapatka, Peter Lichter, Aurélie Ernst

## Abstract

*In vitro* assays for clustered DNA lesions will facilitate the analysis of the mechanisms underlying complex genome rearrangements such as chromothripsis, including the recruitment of repair factors to sites of DNA double-strand breaks. We present a novel method generating localized DNA double-strand breaks using UV-irradiation with photomasks. The size of the damage foci and the spacing between lesions are fully adjustable, making the assay suitable for different cell types and targeted areas. We validated this set-up with genomically stable epithelial cells, normal fibroblasts, pluripotent stem cells and patient-derived primary cultures. Our method does not require a specialized device such as a laser, making it accessible to a broad range of users. Sensitization by BrdU incorporation is not required, which enables analyzing the DNA damage response in post-mitotic cells. Irradiated cells can be cultivated further, followed by time-lapse imaging or used for downstream biochemical analyses, thanks to the high-throughput of the system. Importantly, we showed genome rearrangements in the irradiated cells, providing a proof of principle for the induction of structural variants by localized DNA lesions.

## INTRODUCTION

Protecting the DNA from the most dangerous lesions threatening its integrity, i.e. double-strand breaks (DSBs), is vital for a cell. The complexity of the processes that have evolved to fulfill this task and ensure a faithful DNA damage repair is still insufficiently understood. Recently, catastrophic events characterized by dozens to hundreds of clustered DNA DSBs occurring simultaneously have been described, which in some cases lead to cancer cells^1,2^. During such processes, the genome surveillance mechanisms and repair machinery are overwhelmed, leading to complex genome rearrangements^3,4^. To study the mechanisms underlying localized complex genome rearrangements such as chromothripsis, including the recruitment of repair factors to sites of clustered DSBs, a robust assay to efficiently induce clustered DNA DSBs is required.

Existing protocols, based for instance on the combination of 5-bromo-2-deoxyuridine (BrdU) labeling and ultraviolet C (UVC) irradiation through porous membranes^5^, create localized DNA DSBs. However, the requirement for BrdU incorporation for efficient damage induction limits the applications of such micro-irradiation methods to mitotic cells and prevents long-term follow-up of the irradiated cells. Laser-based assays allow circumventing this restriction^6^, but generally offer a relative low throughput, which does not support downstream biochemical analyses. In addition, laser-based assays necessitate highly specialized equipment and set-ups that are not necessarily technically accessible to any laboratory.

More than 20 years ago, Nelms and colleagues developed a pioneering method to examine DNA repair within intact cells^7^. They used ultrasoft x-rays to induce DNA DSBs in defined subnuclear volumes of human fibroblasts and to visualize DNA repair at such sites. However, this type of partial volume irradiation induces wide stripes of damage within the nuclei, and not localized foci affecting one or a few chromosomes per nucleus. In addition, this type of methods requires an x-ray source.

More recently, chromosomal rearrangements induced by ionizing radiation using a proton microbeam irradiation system were reported and described as “chromothripsis-like” by the authors^8^. Despite the importance of such proof of principle experiments, the choice of a genomically unstable cancer cell line as a model to artificially induce complex chromosome rearrangements lessens the impact of these findings.

We developed an assay to induce sites of DNA DSBs in a high-throughput manner and we applied the method to a range of cell systems.

## MATERIAL AND METHODS

### Induction of clustered DNA DSBs by photomask-based irradiation

To induce DNA lesions, we used high energy UV light with a wavelength of 280 nm emitted by a 20 mW UV LED (Thorlabs), which is collimated by a quarz lens (Thorlabs). LED and lens are mounted on a lens tube system (Thorlabs) and power levels of the LED are adjusted with a manual LED controlling System (Thorlabs). To induce lesions in the order of 1 μm diameter we designed chrome photomasks with arrays of transparent circular areas allowing the transmission of the emitted UV-light. The masks were manufactured by ROSE Fotomasken according to our design. For irradiation experiments, cells are either cultivated on glass slides that are subsequently put on the chrome mask (with the cells pointing towards the chrome mask surface) or seeded directly onto the mask. Irradiation intensities can be regulated in the range of a few hundred μJ/cm^2^ up to several hundred mJ/cm^2^ by adjusting the time of irradiation. After irradiation the cells can be cultivated for further analysis, harvested for biochemical experiments or subsequently studied in live cell microscopy if cells contain labeled DNA-damage reporters. By using a self-written python script (available at https://github.com/ko2001/ct2020) utilizing the open source modules (python3.6, GDSII Library gdspy version 1.3.1) the design of the chrome mask can be easily adapted to specific needs and requirements of a particular experiment (spatial distances and adjustable sizes of the transparent areas).

### Culture conditions

hTERT RPE-1 (ATCC® CRL-4000™) cells (RPE1) were cultivated in DMEM/F12 medium supplemented with 10% fetal calf serum (FCS), 2 mM L-Glutamine, 100 I.U./ml penicillin and 100 μg/ml streptomycin. Induced pluripotent stem cells (iPSCs, line 6-a, 06C53141) were cultured on Matrigel (Corning) pre-coated culture dishes and fed with mTeSR (Stem cell Technologies) supplemented with 100 I.U./ml penicillin and 100 μg/ml streptomycin. Li Fraumeni syndrome fibroblasts (LFS) were cultivated in MEM-alpha supplemented with 1% non-essential Amino acids, 10% FCS, 100 I.U./ml penicillin and 100 μg/ml streptomycin. BJ (ATCC® CRL-2522) cells were cultivated in DMEM medium supplemented with 10% fetal calf serum (FCS), 2 mM L-Glutamine, 100 I.U./ml penicillin and 100 μg/ml streptomycin.

### Serum starvation and mitotic arrest

For serum starvation assays, BJ cells were cultivated in DMEM medium supplemented with 2 mM L-Glutamine, 100 I.U./ml penicillin and 100 μg/ml streptomycin but without FCS. Starvation was performed on exponentially growing cultures for 48 h.

### CRISPR/Cas9 mediated inactivation of *TP53*

In order to generate cell lines that lack functional TP53, two guide RNAs targeting exon 6 and exon 8 of human *TP53* where cloned into the vector system pX330 and electroporated in the target cells utilizing the neon electroporation system (ThermoFisher) (two 20 μs pulses at 1200 V). 24h after electroporation, p53-negative cells were selected utilizing 20 μM Nutlin3a for 5 passages. For iPSCs individual colonies were isolated and expanded. Monoclonal lines were generated using limited dilution and expanded. P53 protein levels where verified using immunoblotting.

### Generation of monoclonal lines

Monoclonal p53-negative RPE1 sublines were generated using limiting dilution on 96 well plates. After seeding, the cells were cultivated till visible colonies were observed. Only wells that contained one single colony were further expanded, and the resulting cell lines were used for subsequent experiments.

### Generation of stable lines expressing 53BP1-eGFP

RPE1 cells co-expressing 53BP1-eGFP and Lifeact-RFP were generated using two individual pIRES-Neo3 vector systems (Clontech) harboring either the coding sequence of 53BP1 (representing amino acid 1220-1711) N-terminally fused to eGFP or the coding sequence of Lifeact C-terminaly fused to RFP. Vectors were co-electroporated into RPE1 *TP53* knockout cells utilizing the Neon electroporation system (ThermoFisher) and selected with 1mg/ml geneticin (Calbiochem). After several passages the cells where FACS-sorted on a FACSAria to isolate double positive cells.

### Whole-genome sequencing

Purified DNA was quantified using the Qubit Broad Range double-stranded DNA assay (Life Technologies) Genomic DNA was sheared using an S2 Ultrasonicator (Covaris). Whole genome sequencing and library preparations were performed according to the manufacturer’s instructions (Illumina). The quality of the libraries was assessed using a Bioanalyzer (Agilent). Sequencing was performed using the Illumina HiSeq 4000 platform.

### Copy number profiling

CNMops (v.1.28.0)^9^ was used for copy number profiling, DNAcopy (v1.56.0) was used for copy number segmentation. Three non-irradiated control clones (control #04, control #09 and control #24) were pooled as controls for the copy number estimation. Copy number profiling was performed for 7 irradiated clones. All lines were sequenced by low-coverage whole-genome sequencing at ∼1.5X coverage. A bin size of 64kb was used for copy-number estimation. Copy-number profiles were visualized by CNMops.

### Quantification of single base substitution (SBS) signatures and single nucleotide variants (SNV) calling

Single nucleotide variants (SNV) were called by Mutect2^10^. Resulting somatic SNVs were converted to 96-classes of trinucleotide context profiles by the PCAWG signature preparation tool^11^. Trinucleotide context profiles were used as input for sigProfiler^11^ to perform a mutational signature analysis and to retrieve the signature exposures of 45 SNV signatures from COSMIC single base substitution (SBS) signatures V3.

(https://cancer.sanger.ac.uk/cosmic/signatures/SBS/)

### Immunofluorescence staining procedure

For immunofluorescence staining, cells were fixed with ice-cold 4% formaldehyde in PBS for 5 min. After fixation, cells were permeabilized in 0.1% Triton-X100 in PBS. Antibodies were used in the following dilutions: Thymine dimer 1:200 (Abcam), γH2AX1:200 (Millipore), 53BP1 1:750 (Santa Cruz), Ki67 1:100 (Abcam). After three washing steps with PBS, incubation with secondary antibodies was performed (Life Technologies). All antibody reactions were carried out in blocking buffer containing 0.2% Triton. After three final washing steps with PBS, the cover slips were rinsed in distilled water, subjected to ethanol for 10 seconds, dried and embedded in DAPI Fluoromount G (southern biotech). Immunofluorescence analysis was performed utilizing a Deltavision microscope and the SoftWorx 4.9 software (Applied precision/GE Healthcare) or an Axio Zeiss Imager.M2 microscope equipped with the ZenSoftware using 63X 1.3 NA oil-immersion objectives.

### Quantification of the intensity of the foci

Quantification of foci intensities was performed with an ImageJ macro adapted from a publicly available macro (https://imagej.nih.gov/ij/source/macros/Circle_Tool.txt). For each individual experiment, at least 200 foci where manually selected and the average mean intensities of the selected areas were recorded. Background evaluation was performed analogously.

### Comet assay and quantification

The alkaline single cell gel electrophoresis assay (comet assay) was performed as previously described^12,13^, with some modifications. RPE1 cells were seeded on comet assay slides (Trevigen) in quadriPERM culture dishes (Sarstedt) with 1×10^5^ cells in 5 ml culture medium per compartment. The cells were allowed to attach and subsequently exposed to high energy UV light (see above). As a positive control, DNA damage was introduced by 5 Gy irradiation using a 137Cs radiation source with a dose rate of 1 Gy min^−1^. Treated cells were coated with 50µl/slide of 0.7% low-melting-temperature agarose (SeaKem). The coating was either done directly after treatment or after allowing 1 or 6 hours’ time for DNA repair at RPE1 cell culture conditions. After coating, the cells were lysed overnight at 4°C. Subsequently, the slides were transferred to a horizontal electrophoresis unit filled with alkaline electrophoresis buffer. After 20 min of DNA unwinding, electrophoresis was performed at 4°C at 0.8 V/cm for 20 min. Slides were neutralized and stained with SYBR Green (Molecular Probes). Fifty-one comets/slide area on three different slides, i.e. a total of 153 cells per data point were selected at random and evaluated by fluorescence microscopy using an imaging software (Kinetic Imaging, Komet 6.0, Andor Technology). The extent of DNA damage was measured quantitatively by the parameter ‘Olive tail moment’. Results are expressed as mean ± standard error.

### Analysis of apoptotic fractions by flow cytometry

Apoptotic cell death was quantified by FITC Annexin V (BD Pharmingen) /7-AAD (BD Pharmingen) staining. Cells were trypsinized using 0.25% trypsin, washed in PBS and centrifuged at 1200 rpm for 5 min. Cells were resuspended in 10% 7AAD with 10% AnnexinV-FITC in 1X Annexin binding buffer. Single stains of 10% 7AAD or 10% Annexin V-FITC were used as controls. For 10% 7AAD single stain control, the cells were heated on 80°C heating block for 10 min to induce cell death. Resuspended cells were incubated at 4°C for 15 min in the dark. 1X Annexin binding buffer was added and cells were immediately analyzed using BD FACS Canto.

### Time-lapse imaging

Time-lapse imaging of stable RPE1 cells expressing 53BP1-eGFP and Lifeact-RFP was performed in a 37°C climate chamber attached to a Deltavision microscope system and the SoftWorx 4.9 software (Applied Precision/GE Healthcare) using a 40x 1.3 NA oil-immersion objective. Cells were seeded on the mask 24h before the irradiation. After irradiation the cells where sealed on the chrome mask in a self-built medium-filled chamber, consisting of a 100 μm thick silicon frame and a #1.5 coverslip protecting the cells from drying out. Images were taken at the indicated time points.

### Metaphase spreads

Mitotic cells were rinsed of the cell culture dish using DPBS. Cell pellets were washed twice with PBS at room temperature and gently pelleted at 800xg. Washed cells were incubated in hypotonic swelling buffer (0.56% KCl) for 6 min at 37°C, pelleted and fixed with 3 parts methanol and 1 part glacial acetic acid. Fixed cells were either dropped on pre-cleaned glass slides or stored at −20°C. For microscopic analysis mitotic spreads were either sealed with DAPI Fluoromount G, subjected to M-FISH or Chromosome 14 painting.

### M-FISH

Seven pools of flow-sorted human chromosome painting probes were amplified and directly labeled using seven different fluorochromes (DEAC, FITC, Cy3, Cy3.5, Cy5, Cy5.5 and Cy7) using degenerative oligonucleotide primed PCR (DOP-PCR). Metaphase chromosomes immobilized on glass slides were denatured in 70% formamide/2xSSC pH 7.0 at 72°C for 2 minutes followed by dehydration in a degraded ethanol series. Hybridization mixture containing labeled painting probes, an excess of unlabeled cot1 DNA, 50% formamide, 2xSSC, and 15% dextran sulfate were denatured for 7 minutes at 75°C, pre-annealed at 37°C for 20 minutes and hybridized at 37°C to the denatured metaphase preparations. After 48 hours the slides were washed in 2xSSC at room temperature for 3x 5 minutes followed by two washes in 0.2xSSC/0.2% Tween-20 at 56°C for 7 minutes, each. Metaphase spreads were counterstained with 4.6-diamidino-2-phenylindole (DAPI) and covered with antifade solution. Metaphase spreads were recorded using a DM RXA epifluorescence microscope (Leica Microsystems) equipped with a Sensys CCD camera FISH software and images were processed on the basis of the Leica MCK software and presented as multicolor karyograms (Leica Microsystems Imaging solutions)

### FISH – Chromosome 14 painting probes

Whole chromosome paint probe (Metasystems) was applied on the slides with metaphase spreads and covered and sealed with coverslips and rubber cement. Denaturation was done at 78°C for 4 min and the slides were incubated overnight in a humidified chamber at room temperature for hybridization. Coverslips and rubber cement were removed the next day and post-hybridization washes were performed with 0.4 SSC (pH 7.0) at 72°C for 8 min and 2xSSC, 0.05% Tween-20 (pH 7.0) at room temperature for 5 min. Slides were rinsed in ddH20 and air-dried. DAPI Fluoromount-G (Southern Biotech) was used for counterstaining the nuclei. Images were acquired using an Axio Zeiss Imager.M2 microscope. All images were captured using a 63X 1.3 NA magnification. A minimum of 60 nuclei was counted per sample.

### Immunoprecipitation and Immunoblotting

Cell lysates were generated using RIPA buffer (10 mM Tris-HCl at ph 8.0, 1 mM EDTA, 1% Triton X-100, 0.5% sodium deoxycholate, 0.1% sodium dodecyl sulfate, 150 mM NaCl) containing protease inhibitors (Roche complete) and 100U/ml benzonase (Merck). Immunoprecipitation (IP) was performed with protein A-coupled magnetic beads (Invitrogen Dynal). 20 μg yH2AX antibody, raised in rabbit (Cell signaling), was incubated with 30 μl of bead-slurry for 15 min at 4°C. IP was carried out for 30 min with 0.5 ml lysate incubated with the coupled bead-slurry at 4°C under constant rotation. Washing was done with lysis buffer, three times 5 min with gentle agitation at 4°C. Elution of precipitates was performed with Laemmli sample buffer (2% SDS, 10% glycerol, 75 mM Tris-HCl pH 6.8, 50 mM dithiothreitol) for 5 min at 95°C. Lysates and IP eluates were separated on 4-12% gradient gels (Invitrogen) and blotted on nitrocellulose membranes (Amersham, GE Healthcare) using a semi-dry blotter and a 10 mM CAPSO buffer system adjusted at pH 11. Membranes were blocked with 5% milk powder at room temperature for 1h. Detection of yH2AX was performed with an anti-yH2AX antibody, raised in mouse (Millipore), according to the manufacturer’s protocol. All antibody reactions were carried out in 50 mM Tris-HCl, pH 8.0, 150 mM NaCl and 0.05% Tween 20 with 5% milk powder at room temperature for 1h.

### T4EndoV assay

Purified PX330 (Addgene) vector DNA was fully digested with EcoRV (New England Biolabs) and treated with the UV irradiation system without a photomask. Irradiated and control DNA was either loaded directly on a 1% agarose gel or incubated with T4 endonuclease V (New England Biolabs) for 30 minutes at 37°C and subsequently analyzed by agarose gel electrophoresis using a 1% agarose gel.

### Cell cycle analysis by flow cytometry

Cell cycle analysis was conducted using propidium iodide (Sigma Aldrich). Treated and untreated cells were harvested and resuspended in 70% ethanol at 4°C for 2h followed by centrifugation at 1200 rpm for 5 min. Ethanol supernatant was discarded, cell pellet was resuspended in propidium iodide solution (1mg/ml) and incubated at 4°C for 30 min. FACS measurement was done with BD FACS Canto.

## RESULTS

### Efficient generation of localized DNA double-strand breaks (DSBs) in a highly versatile manner with fully adjustable irradiation areas

We developed a system allowing a tight control of the exact area of DNA damage and of the number of foci per nucleus. By designing customized chrome photomasks, we reproducibly induce DSB foci of user-defined sizes and spatial arrangements (Figure 1 a). With this system, nuclear areas down to 1 μm in diameter can be irradiated. Irradiation of chromosomally stable near-diploid Retinal Pigment Epithelial (RPE1) cells showed a highly efficient induction of clustered DSBs in interphase and metaphase cells, as visualized by the co-localization of UV-induced thymine dimers with DNA DSB markers γH2AX and 53BP1 (Figure 1 b-f). Importantly, the cells are placed directly onto the photomask and can be either fixed after the irradiation or kept in culture for downstream analyses.

**Figure 1.**
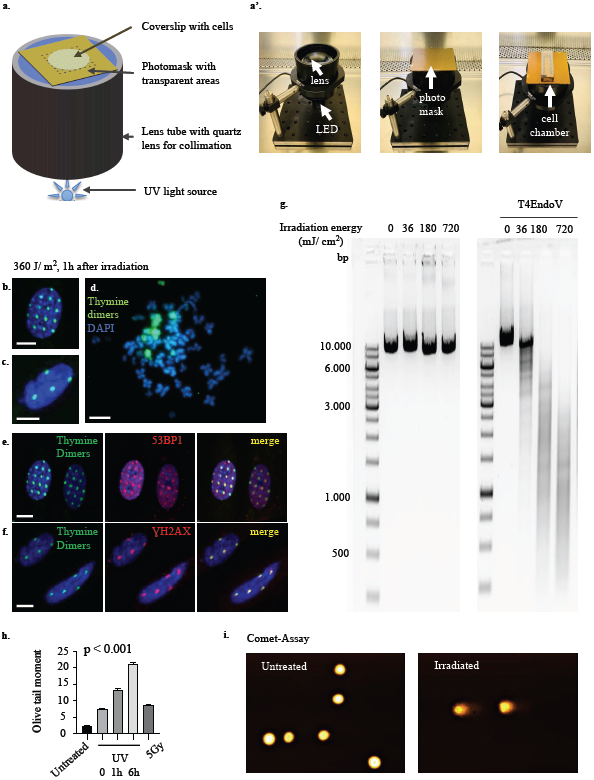
**a**, Scheme and photographs of the set-up for the generation of localized DNA double-strand breaks by UV light in combination with a photomask. **b-d**, Immunofluorescence analyses of UV-induced thymine dimers in wild-type RPE1 cells after UV irradiation through masks of different designs (transparent areas with 1 μm diameter, 5 μm spacing in b; transparent areas with 2 μm diameter, 10 μm spacing in c). **d**, Thymine dimer staining on metaphase spreads from irradiated cells shows that a subset of chromosomes was affected by the induced damage. **e-f**, UV-induced thymine dimers co-localize with DNA DSB markers 53BP1 and γH2AX. Scale bar, 10 μm. Images are representative of three independent experiments. **g**, The linearized pX330 plasmid was irradiated with 200 μW/cm^2^ UV light or mock treated (no UV, time 0). Without treatment with the endonuclease T4EndoV, we did not observe any major change in the size of the linearized plasmid, even after 60 minutes of irradiation (left panel). However, we detected a shift towards smaller sizes and a smear on the gel after T4EndoV treatment, already visible after irradiation with 36 mJ/cm^2^ (right panel, +UV, +T4endoV). Gel pictures are representative of two independent experiments. **h**, Determination of DNA damage induced by irradiation with the alkaline single-cell electrophoresis assay. Wild-type RPE1 cells were irradiated with UV (100 J) and the DNA damage induced was analyzed immediately after irradiation, 1 hour and 6 hours after irradiation. Irradiation with γ-rays (5 Gγ) was used as a positive control (sample taken directly after irradiation). Each bar shows the parameter ‘Olive tail moment’ (mean ± SE) determined on 51 randomly selected comets per slide and three different slides (n=153) from two independent experiments. Statistical significance was tested using Kruskal-Wallis tests with Dunn’s multiple comparison adjustments. All comparisons were highly significant (p<0.001) except for the comparison between the positive control (5 Gy) and the UV irradiation (both directly after irradiation). **i**, Representative Comet assay micrographs for untreated cells and for cells irradiated with 5 Gy (positive control).

In order to assess the proportion of DSBs induced as a secondary effect due to the thymine dimers (as opposed to the less likely direct induction of DSBs by UV light), we used the endonuclease T4EndoV. This repair enzyme recognizes UV-induced dimers and cleaves the DNA at the lesion sites^14,15^. We first linearized the pX330 plasmid with EcoRV and then irradiated the plasmid with 36, 180 or 720 mJ/cm^2^. Without treatment with T4EndoV, we did not observe any major change in the size of the linearized plasmid, even after applying energies as high as 720 mJ/cm^2^ (Figure 1 g, left panel). However, we detected a shift towards smaller sizes and a smear on the gel after T4EndoV treatment, already visible after the application of 36mJ/cm^2^ (Figure 1 g, right panel). This finding suggests that the vast majority of the DSBs detected by the γH2AX and 53BP1 antibodies (Figure 1 e-f) originate from secondary enzymatic reactions at the lesion sites, rather than from directly induced DSBs from the UV light itself.

To quantify the DNA breaks generated by the irradiation and to compare the DNA damage levels with those induced by ionizing radiation, we performed COMET assays (Figure 1 h-i). Irradiation in combination with the UV photomask leads to lesions close to the detection limit of the COMET assay (as only a small proportion of the nucleus is irradiated). Therefore, we analyzed irradiated cells without mask as a proof of concept for the type of damage induced by this irradiation system. Immediately after UV irradiation without photomask, the DNA damage levels were in the same range as when applying 5 Gy of ionizing radiation (no significant difference to the positive control, Kuskal-Wallis test). Six hours after UV irradiation, the DNA damage levels kept increasing, indicating secondary lesions.

We next performed a dose-response experiment to quantify the intensity of the damage foci with increasing irradiation energies. We showed a nearly-proportional increase of the signal intensity for the DNA DSB marker γH2AX when applying irradiation energies of 100, 200 and 300 J, respectively (Figure 2 a). To analyze the kinetics of the DNA damage response, we quantified the intensity of the damage foci starting from 30 minutes after irradiation and up to 24 hours later (Figure 2 b). The intensity of the foci was still increasing 30 minutes after the irradiation but decreasing six hours after irradiation, in agreement with the time course revealed by the COMET assay. After 24 hours, the DNA damage levels were not yet back to background levels, as shown by residual γH2AX foci.

**Figure 2.**
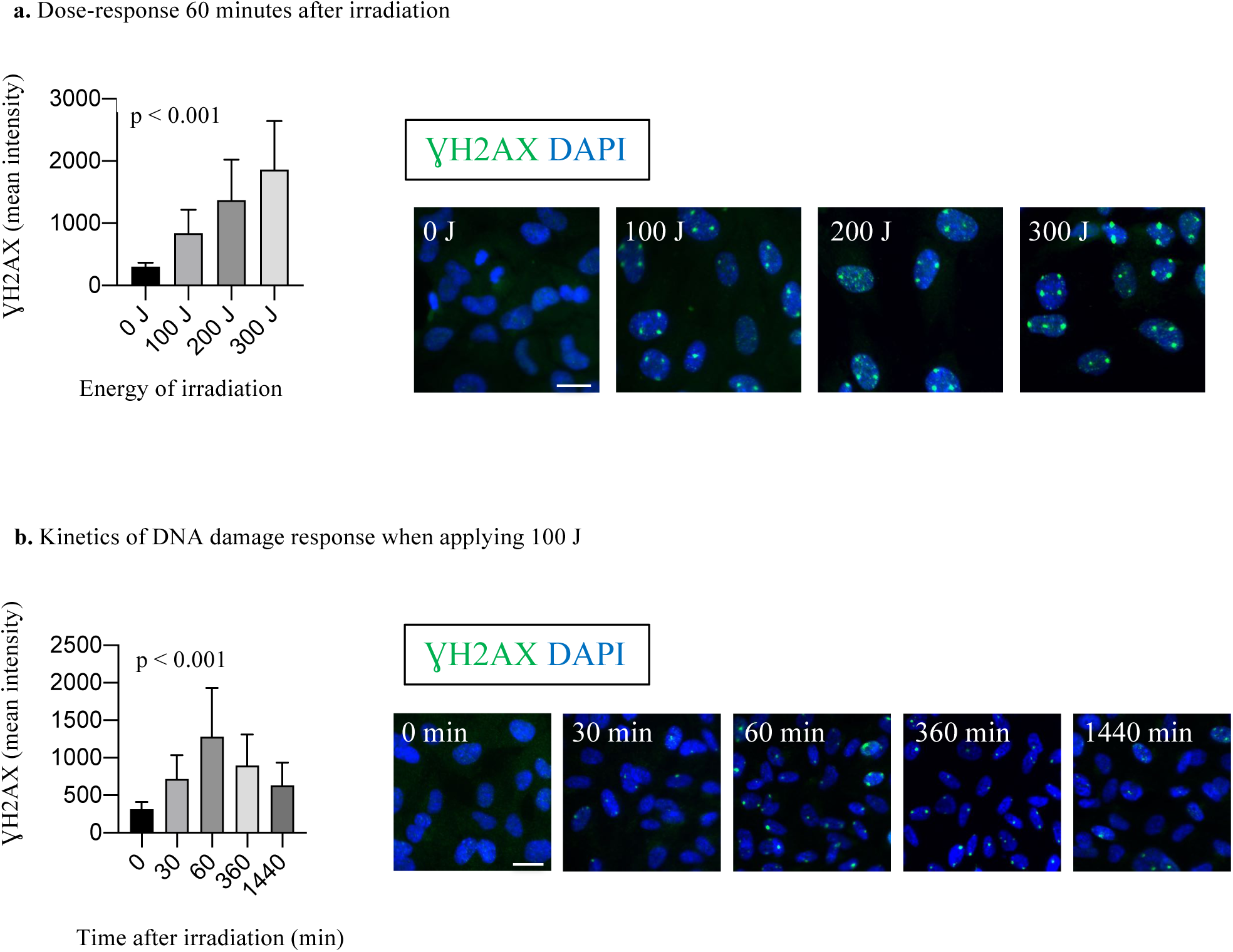
Dose dependency and time-course analysis. Immunofluorescence analyses of UV-induced DNA DSBs in wild-type RPE1 cells after UV irradiation in combination with a photomask (transparent areas with 2 μm diameter, 10 μm spacing). The fluorescence intensity of the γH2AX foci was quantified. **a**, The average signal intensity of the γH2Ax foci is shown for each energy of irradiation applied. Cells were fixed 60 minutes after irradiation. **b**, The average signal intensity of the γH2AX foci is shown as a time-course starting from the time point of the irradiation. All cells were irradiated with 100 J. Bar graphs show average values and standard deviation for three independent experiments in a and two independent experiments in b. The untreated controls (0J in a and t=0 in b) show background fluorescence values. Scale bar, 20 μm. Statistical significance was tested using Kruskal-Wallis tests with Dunn’s multiple comparison adjustments. All comparisons were highly significant (p<0.001).

BrdU can be used to sensitize cells to UV damage and increase the size of the DSB foci (Supplementary Figure 1) ^5^. However, the addition of BrdU is not a requirement for our set-up, as our system leads to highly efficient localized DSB induction even without sensitization with BrdU (Supplementary Figure 1). Therefore, we can efficiently target non-dividing cells and avoid potential side-effects associated with BrdU treatment^16^. As a proof of concept, we induced mitotic arrest in normal fibroblasts (BJ cells) by serum starvation. We then irradiated non-mitotic starved cells and control non-starved cells and stained for γH2AX as well as for proliferation marker Ki67 (Supplementary Figure 2). Importantly, serum-starved cells were negative for proliferation marker Ki67, as expected, but positive for DNA damage marker γH2AX.

We next tracked DSB foci and monitored the morphology of the cells after the damage induction. For this, we generated a p53-deficient RPE1 line stably expressing eGFP-53BP1 and LifeAct-RFP to visualize DSBs and actin, respectively. Induction of DSB lesions in these cells and live-cell imaging for 24h after damage showed that irradiated cells are able to complete mitosis (Figure 3 a-b). Therefore, our system allows following irradiated cells over time and analyzing the evolution of the induced lesions in damaged cells.

**Figure 3.**
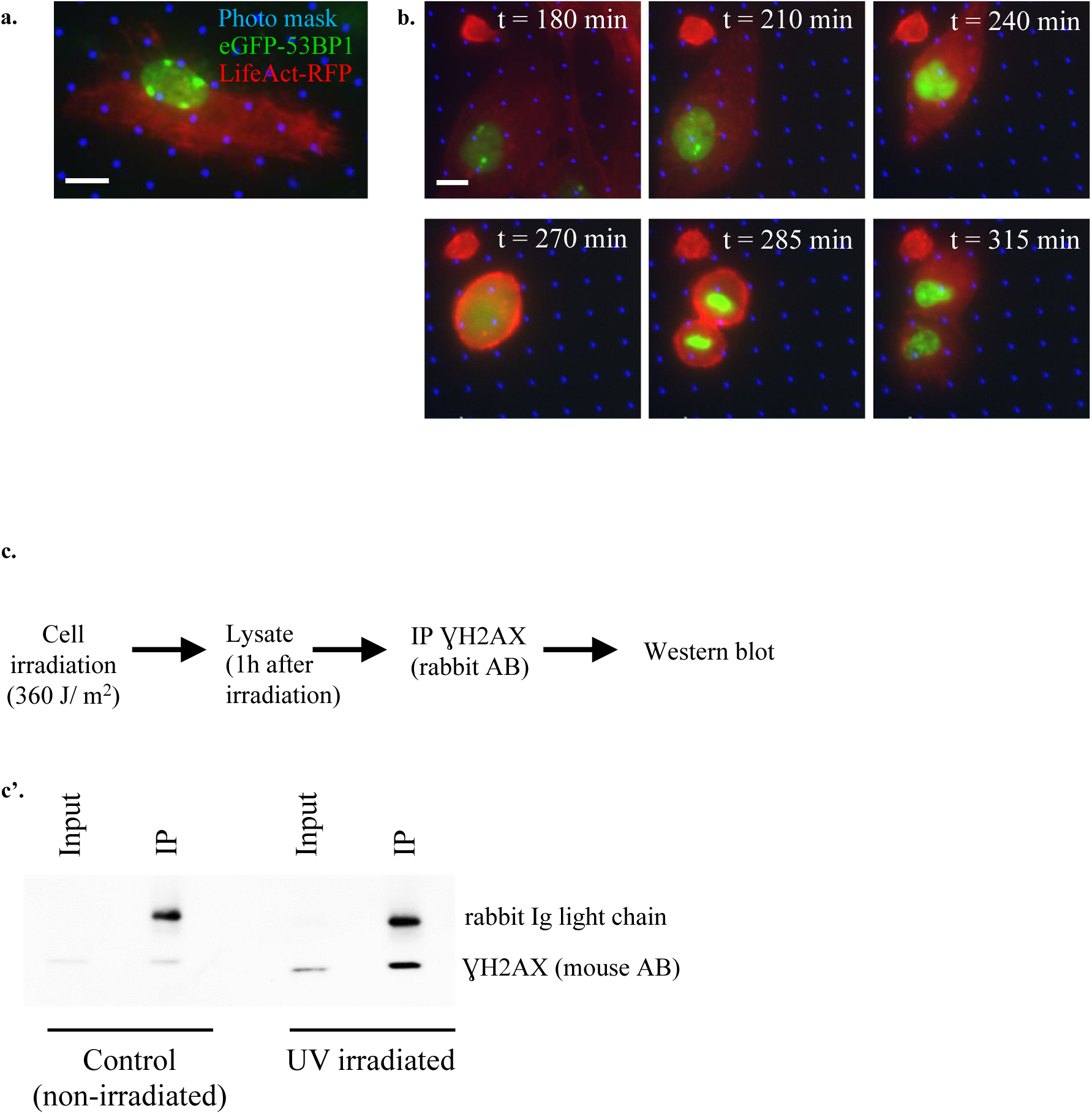
Downstream applications for efficient generation of localized double-strand breaks: live-cell imaging of DNA damage repair and biochemical analysis of the foci components. **a, b**, P53-deficient RPE1 cells (whereby *TP53* was inactivated by CRISPR/Cas9) stably expressing eGFP-53BP1 to monitor 53BP1 at DNA DSBs and LifeAct-RFP to follow cell morphology were grown on the mask and tracked by time-lapse imaging for 18 h after the damage induction (t indicates the time in minutes after irradiation). Blue dots show the position of the transparent areas of the photomask through which the radiation is transmitted. Scale bar, 10 μm. Images are representative of two independent time-lapse experiments. **c**, RPE1 cells were irradiated in combination with the photomask and cell lysates were prepared. Immunoprecipitation and immunoblotting were performed using antibodies against γH2AX. Immunoprecipitation and immunoblotting of non-irradiated cells were done as a control. Images are representative of two independent experiments.

In contrast to most current methods to induce localized DSBs, our approach offers a sufficient throughput for the downstream biochemical analysis of the foci components. To validate this type of applications, we irradiated RPE1 cells and collected cell lysates for immunoprecipitation and immunoblotting with γH2AX antibodies (Figure 3 c). This proof of principle experiment shows that our set-up can be used for the biochemical characterization of the factors present at the damage foci, to identify novel repair factors.

### Phenotypic effects in cells surviving localized DNA DSBs

We then analysed the phenotypic consequences of the damage induction in different cell types. After irradiation of induced pluripotent stem cells (iPSCs) from healthy donors with wild-type p53, most cells die (Figure 4 a, left panel). When the same irradiation dose is applied to iPSCs of the same donor after CRISPR-Cas9-mediated *TP53* knock-out or to iPSCs derived from fibroblasts of patients with Li Fraumeni syndrome (germline mutation in *TP53*), a substantial fraction of the cells survives the irradiation. Immunofluorescence analysis of DSB markers γH2AX and 53BP1 24h after irradiation showed damage foci in the surviving cells (Figure 4 a, lower panels). Likely due to impaired p53 function, *TP53* knock-out iPSCs and Li Fraumeni syndrome iPSCs survive the induced damage. The capacity of such cells to overcome cell cycle arrest and apoptosis triggered by localized DSBs possibly contributes to the frequent occurrence of chromothripsis in cells of patients with Li Fraumeni syndrome^1^.

**Figure 4.**
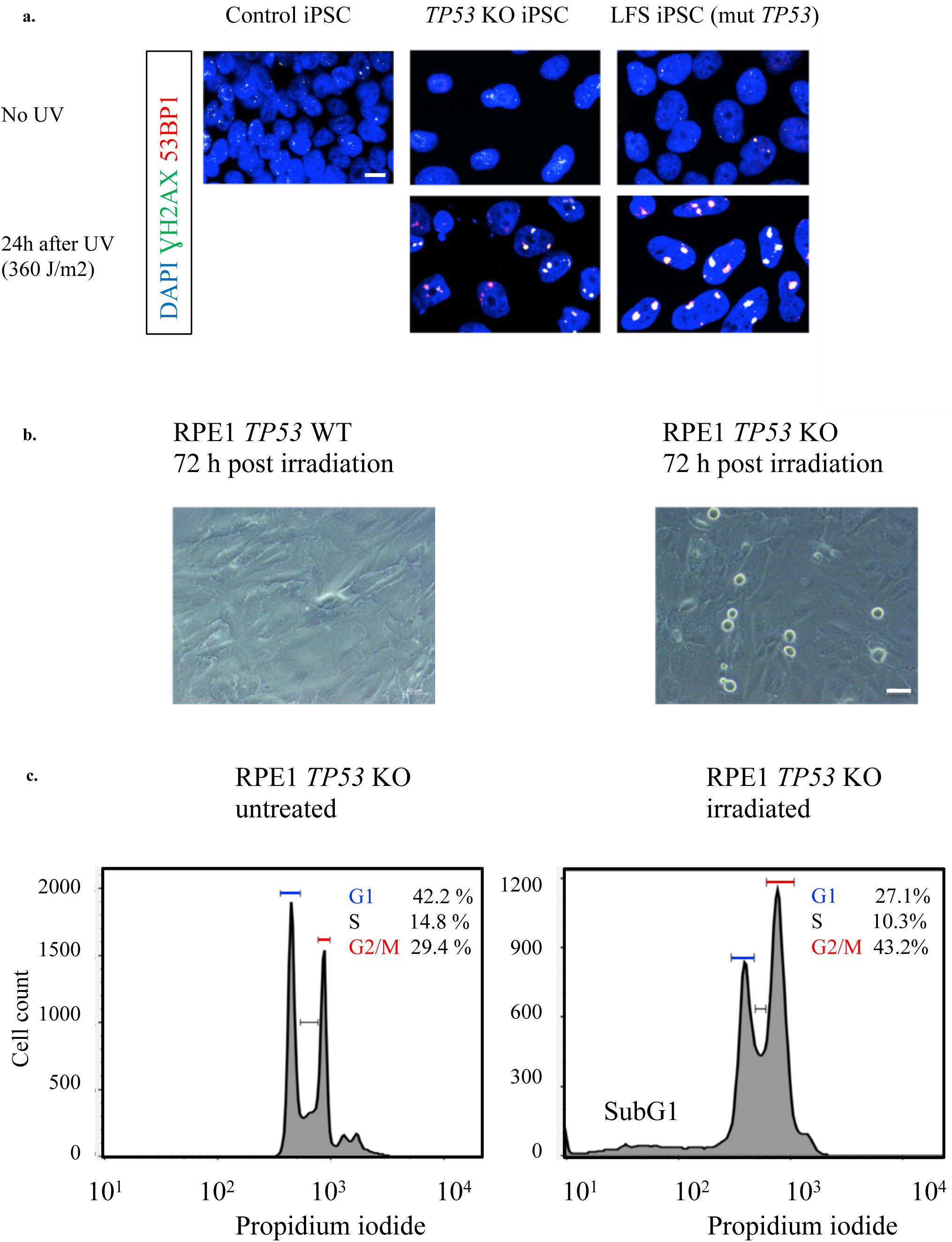
Phenotypic effects of localized DSBs on *TP53* wild-type and *TP53*-deficient iPSCs and RPE1 cells. **a**, Immunofluorescence analysis of DSB markers before and after irradiation in healthy donor iPSCs, *TP53* knock-out iPSCS and iPSCs from a patient with Li Fraumeni (LFS) syndrome, respectively. Healthy donor iPSCs do not survive the irradiation, whereas p53-compromised iPSCs tolerate irradiation and show foci at the sites of damage. Scale bar, 10 μm. Images are representative of three independent experiments. **b**, The mitotic index of *TP53* knock-out RPE1 cells increases after irradiation. Brightfield images are representative of three independent experiments. Scale bar, 50 μm. **c**, Cell cycle analysis of p53-deficent RPE1 cells before and after irradiation, respectively. Flow cytometry plots are representative of three independent experiments.

In RPE1 cells, we observed an increased number of mitotic cells after irradiation in the *TP53*-knock-out line, but not in the irradiated *TP53* wild-type cells (Figure 4 b). In line with this finding, we detected a substantial change in the cell cycle distribution in *TP53*-knock-out cells after irradiation (Figure 4 c), with a 30% increase in the proportion of cells in the G2/M phase of the cell cycle in the irradiated cells. The proportion of apoptotic cells, as shown by the minor SubG1 fraction (Figure 4 c, right panel) and the small proportion of Annexin-positive cells (<5%, Supplementary Figure 3 a) was not significantly increased in RPE1 cells after irradiation. Therefore, the vast majority of RPE1 cells survive the irradiation when applied in combination with the photomask.

Altogether, these findings suggest that in the context of compromised p53 function, cells surviving the induced localized DSBs may acquire an advantage in terms of proliferation. Next, we asked whether the phenotypic effects subsequent to irradiation might go along with chromosome rearrangements induced by the localized DSBs.

### Rearrangements induced by localized DSBs

To assess whether irradiated cells show structural or numerical aberrations, we first did M-FISH analysis of p53-deficient RPE1 cells with and without focal irradiation, respectively. In addition to the rare rearrangements already present in the non-irradiated mother line and reported in the literature, such as a derivative X-chromosome with additional chromosomal material at the terminal end of the q-arm^17^, we detected translocations and aneuploidies (Supplementary Figure 4 a-b). Importantly, we detected more numerical and structural aberrations (five-fold more translocations, two-fold more overall aberrations) in metaphases from the irradiated cells as compared to the non-irradiated cells (Supplementary Figure 4 b). Therefore, the induction of localized DSBs does lead to rearrangements in the irradiated cells, and the detected changes are not merely due to chromosome aberrations accumulated over passaging.

We confirmed these rearrangements in a larger number of metaphase spreads (>60 for untreated and irradiated cells, respectively) using chromosome 14 painting probes (Supplementary Figure 4 c-d). The vast majority (>97% of the analysed metaphases) of *TP53* knock-out non-irradiated metaphase spreads displayed two copies of chromosome 14. However, in the irradiated cells, more than 20% of the metaphase spreads presented structural or numerical aberrations affecting chromosome 14. The most frequent aberrations were aneuploidy (three and more copies of chromosome 14) and translocations, but we also detected one cell with chromosome 14 included in a micronucleus. Interestingly, this type of nuclear structure is tightly linked with DNA damage and with complex genome rearrangements such as chromothripsis^18^.

In addition, we analyzed metaphase spreads of RPE1 *TP53*-knock-out cells that showed increased numbers of mitotic cells 48h after irradiation. Remarkably, we observed fragmented or pulverized chromosomes (Supplementary Figure 4 e), which supports our hypothesis that the vast majority of the observed genomic aberrations are caused by the induction of localized DSBs and not by prolonged passaging.

To analyze copy-number changes and point mutations induced by the irradiation, we performed whole-genome sequencing of 11 single RPE1 *TP53*-knock-out clones (n=4 non-irradiated control clones and 7 irradiated clones). Three of the four non-irradiated clones showed no copy-number aberration (Figure 5, Supplementary Figure 5). One control clone showed a copy-number change on chromosome 19, which was most likely the result of spontaneous copy-number changes over passaging, as described by others^19^. Importantly, four out of seven irradiated clones showed copy-number changes affecting one or two chromosome(s) each (Figure 5, Supplementary Figure 5). Furthermore, irradiated clones showed significantly higher loads of single nucleotide variants, mutational signatures of UV-induced damage and APOBEC activity (Figure 5 e), suggesting APOBEC editing of the fragmented DNA. Interestingly, APOBEC activity was previously linked with complex genome rearrangements such as chromothripsis^20^. These findings, together with the higher proportion of copy-number alterations in irradiated clones, demonstrate that our irradiation system can be used to investigate the mechanisms underlying the induction of clustered DNA DSBs.

**Figure 5.**
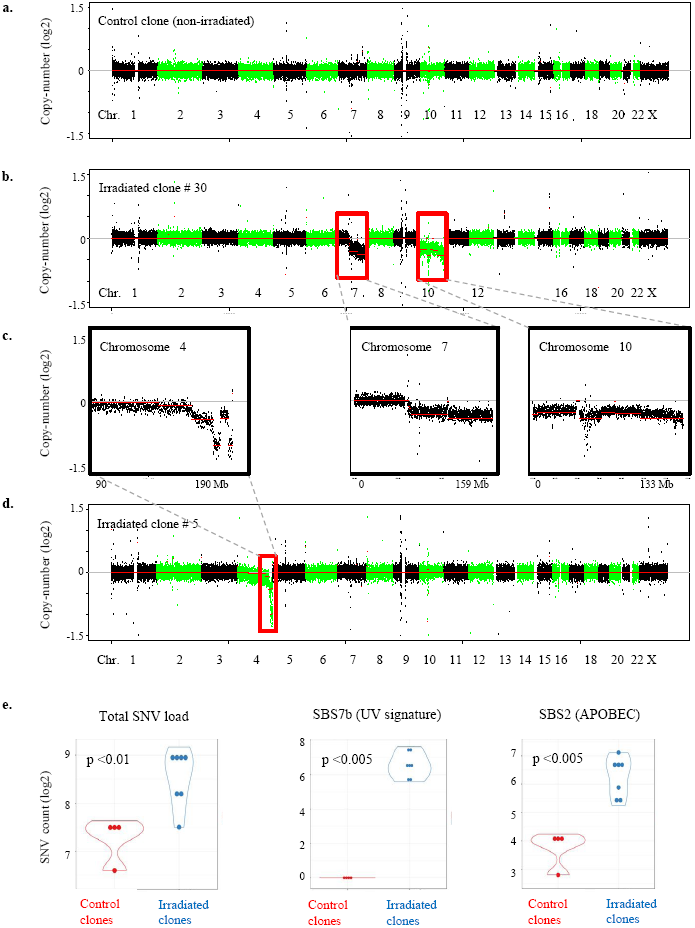
Genome rearrangements induced by localized DNA double-strand breaks. Whole genome sequencing analysis of single clones after irradiation of RPE1 *TP53* KO cells with 300 J shows copy-number alterations **(a-d)** and point mutations **(e)**. Whole-genome views are shown in a, b and d. Individual chromosomes are shown in c. **e**, The total SNV load is shown for each of the four non-irradiated (red) and seven irradiated clones (blue) in the left panel. SNVs linked with mutational signatures of UV light (COSMIC signature SBS7b) and APOBEC activity (SBS2) are shown in the middle and right panels, respectively. Each dot represents one individual clone. Single clones were picked 72 hours after irradiation, expanded and then randomly picked for whole-genome sequencing. Statistical significance was tested using unpaired t-tests (two-tailed).

## DISCUSSION

We developed a method to generate localized DNA DSBs in single cells. In comparison to other micro-irradiation technologies, our system has several unique features. First, due to the regular distribution of the transparent areas through which the damage-inducing beam is transmitted, it allows to easily discriminate real damage foci from false positive signals, which provides a substantial advantage compared to irradiation experiments with micropore membranes that display a random distribution of pores. Second, compared with laser-based micro-irradiation systems that need complex hard- and software setups, our system is relatively straightforward to use and inexpensive, making it accessible to a broad range of users. Third, the throughput of our system is several orders of magnitude higher, as compared with laser-based micro-irradiation systems. While microscopy-based laser systems can sequentially irradiate a maximum of several 100 cells per hour, our method is capable of irradiating 10^6^-10^7^ cells within minutes. Therefore, the large number of cells that can be targeted also enables downstream biochemical experiments (such as co-immunoprecipitation or chip sequencing) requiring amounts of biological material that cannot be obtained with low-throughput systems. Fourth, live cell microscopy can be performed after the damage induction, when the irradiated cells express fluorescently tagged marker proteins and these cells are growing on the photomask. Finally, recording the transmitted light that passes the pores provides an internal reference that can be correlated with the visualized DNA lesions. Thus, we developed a robust tool to induce localized DSBs, which can be applied to a range of cell systems.

A limitation of our method is the requirement for sensitive handling of the cells growing on the mask, as they can detach easily if the experiment is not carried out with sufficient caution. The mask can be designed according to specific needs, with the files used to produce the masks for the current study being available (see Methods) and easily modified for user-defined applications. Furthermore, companies producing photomasks offer services for the generation of individual designs.

Our assay provides unique possibilities to analyze the recruitment of DNA repair factors and the effects of localized DNA DSBs. The availability of *in vitro* model systems to efficiently generate clusters of DSBs is critical for the understanding of complex genome rearrangements. Current methods, such as doxorubicin-induced DSBs^21^, induce genome-wide DNA damage, as compared to the irradiation of a few chromosomes achieved with our system. CRISPR-based approaches, which could potentially be used to introduce localized DSBs, require prior selection of the target sequence. Therefore, nuclease-based strategies offer less flexibility as compared to our method, which allows to restrict the area of damage while potentially affecting any genomic region.

Thus, our approach provides a tool with a broad range of applications in the fields of DNA damage and repair as well as genome rearrangements in cancer cells.

## SUPPLEMENTARY INFORMATION

Supplementary Data are available online.

## ACKNOWLEDGEMENT

We thank Brigitte Schoell and Reinhard Gliniorz for excellent technical support for the M-FISH analysis and for the COMET assays, respectively. We thank Thanos Halazonetis for kindly sharing the 53BP1-eGFP plasmid.

---

**Supplementary Figure 1.**
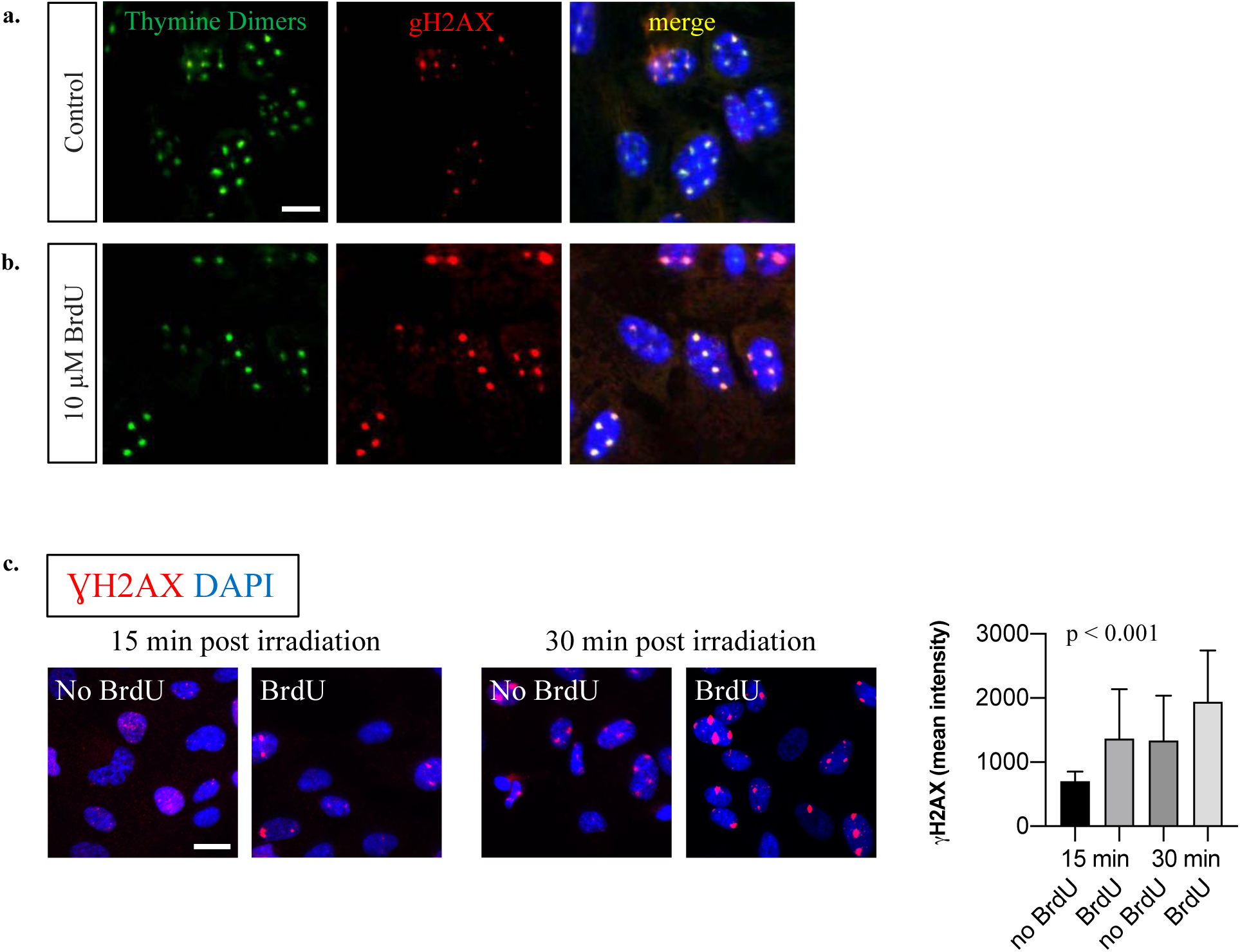
Immunofluorescence analyses of UV-induced thymine dimers and DNA DSBs in RPE1 cells after UV irradiation without **(a)** or with **(b)** BrdU pre-treatment, respectively. BrdU incorporation leads to larger lesions as visualized by the stronger γH2AX signals (middle panels) but is not a requirement for efficient DSB induction. Scale bar, 10 μm. Images are representative of two independent experiments. The irradiation was applied using 600 J/m^2^. Cells where fixed 1h after irradiation. **c**, The mean signal intensity of the γH2AX foci is shown without or with 10 μM BrdU pre-treatment, respectively. All cells were irradiated with 100 J. BrdU incorporation leads to larger lesions as visualized by the stronger γH2AX signals but is not a requirement for efficient DSB induction. Scale bar, 10 μm. Images are representative of two independent experiments. Statistical significance was tested using Kruskal-Wallis tests with Dunn’s multiple comparison adjustments. All comparisons were highly significant (p<0.001) except between BrdU at 15 minutes and no BrdU at 30 minutes.

**Supplementary Figure 2.**
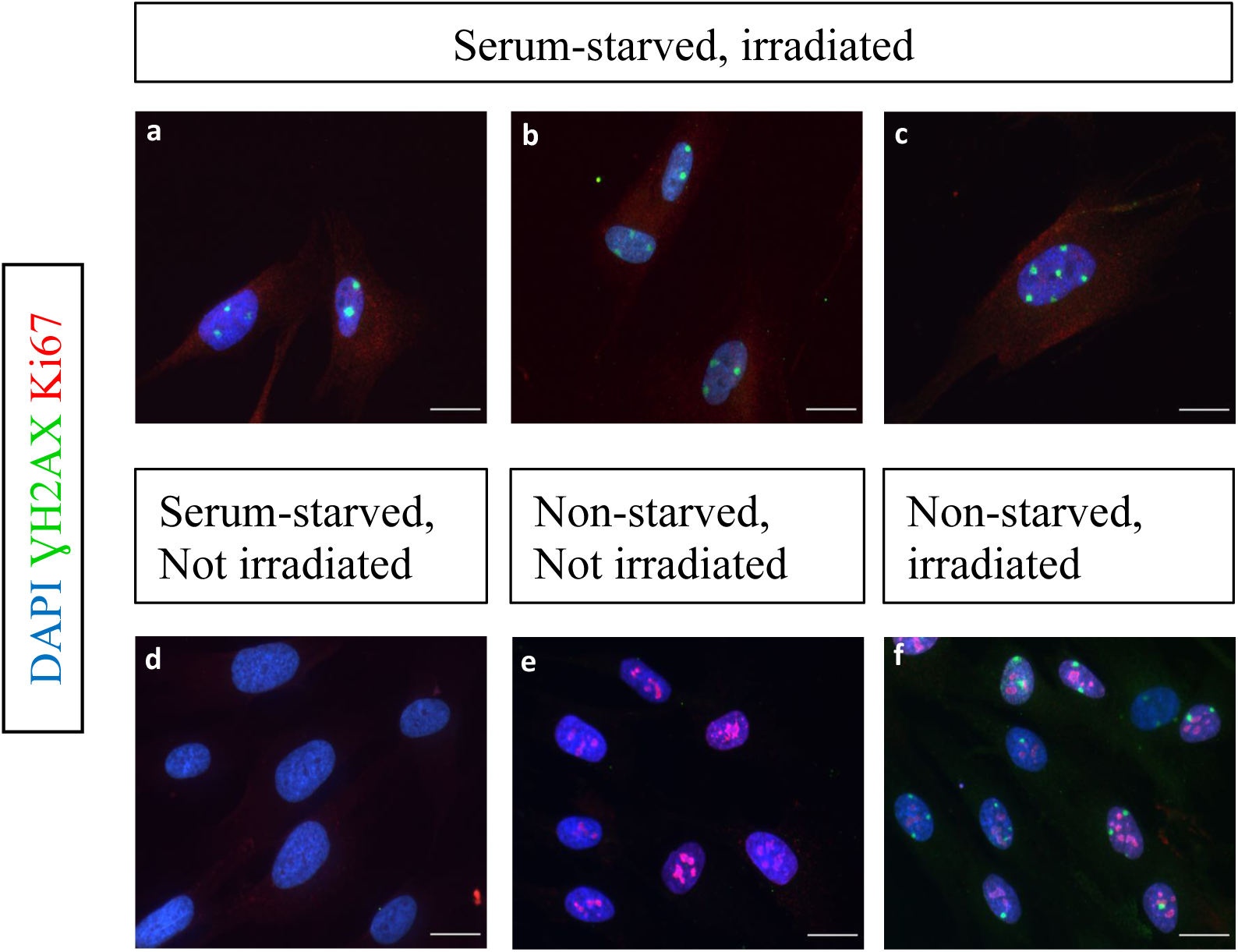
Induction of clustered DNA DSBs in non-mitotic cells. Cell cycle arrest was induced in genomically stable BJ fibroblasts by serum starvation. Immunofluorescence analyses of UV-induced DNA DSBs (γH2AX, green) and proliferation marker Ki67 (red) were performed after irradiation. **a-c**, serum-starved cells 60 minutes after irradiation with 100 J/m^2^. **d**, serum-starved cells without prior irradiation. **e**, non-serum-starved cells without prior irradiation. **f**, non-serum-starved cells 60 minutes after irradiation with 100 J/m^2^. Scalebars, 20 μm.

**Supplementary Figure 3.**
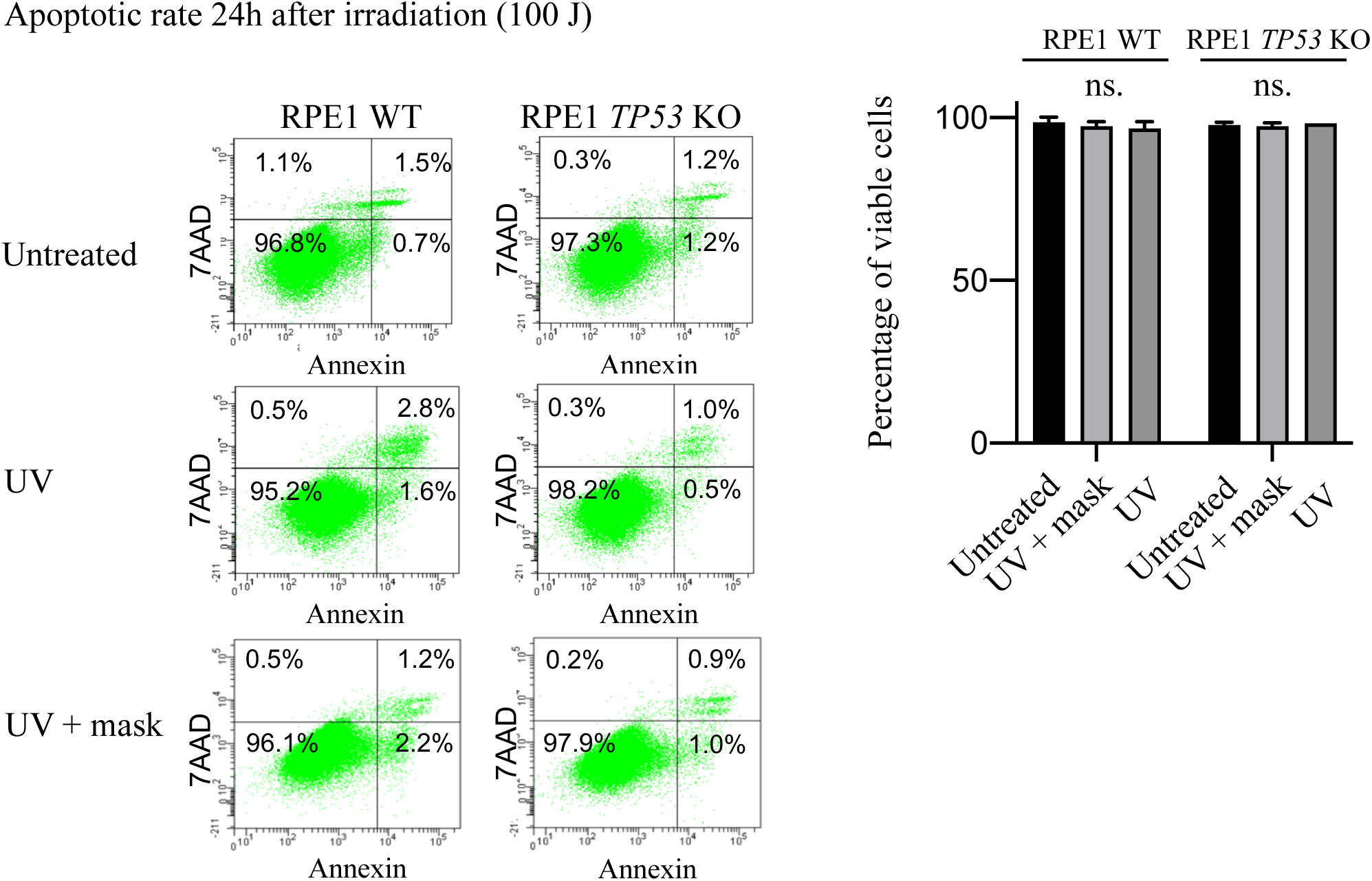
Cell viability after induction of localized damage. Representative FACS analyses of Annexin-V/7AAD stainings and quantifications for three independent experiments. Annexin-V positive cells are apoptotic cells, 7-AAD-positive cells are dead cells.

**Supplementary Figure 4.**
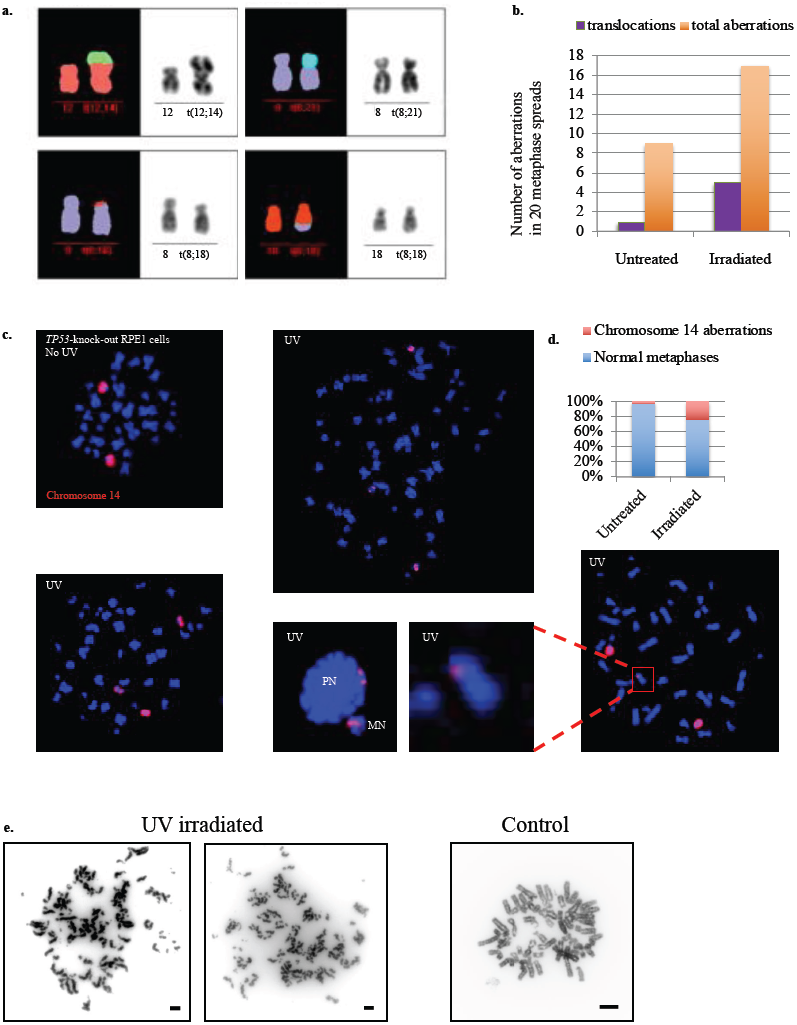
Rearrangements induced by localized DSBs. **a**, Irradiation leads to structural and numerical aberrations, as shown by M-FISH analysis of p53-deficient RPE1 cells before and after irradiation, respectively. Examples of translocations detected in p53-deficient RPE1 cells after irradiation are shown. **b**, Quantification of the structural and numerical aberrations detected by M-FISH analyses in untreated and irradiated p53-deficient RPE1 cells, respectively. **c**, Painting probes for chromosome 14 were hybridized on metaphase spreads of p53-deficient RPE1 cells before and after irradiation, respectively. PN, primary nucleus; MN, micronucleus. **d**, The bar graph shows the percentage of aberrations (numerical and structural) on chromosome 14 detected by counting a minimum of 60 cells for each condition. **e**, Metaphase spreads of RPE1 *TP53*-KO cells 48h after irradiation showing fragmented chromosomes.

**Supplementary Figure 5.**
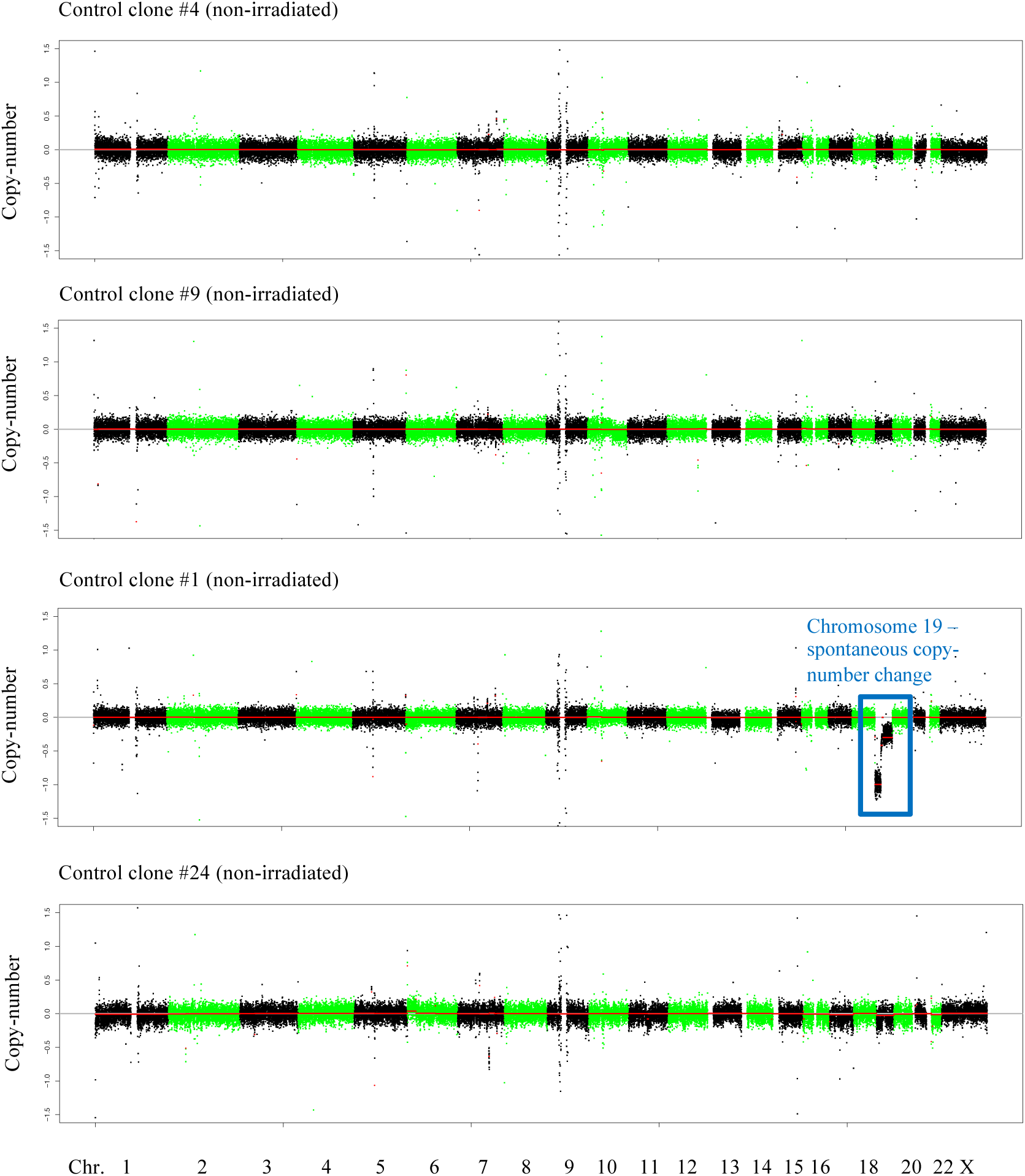
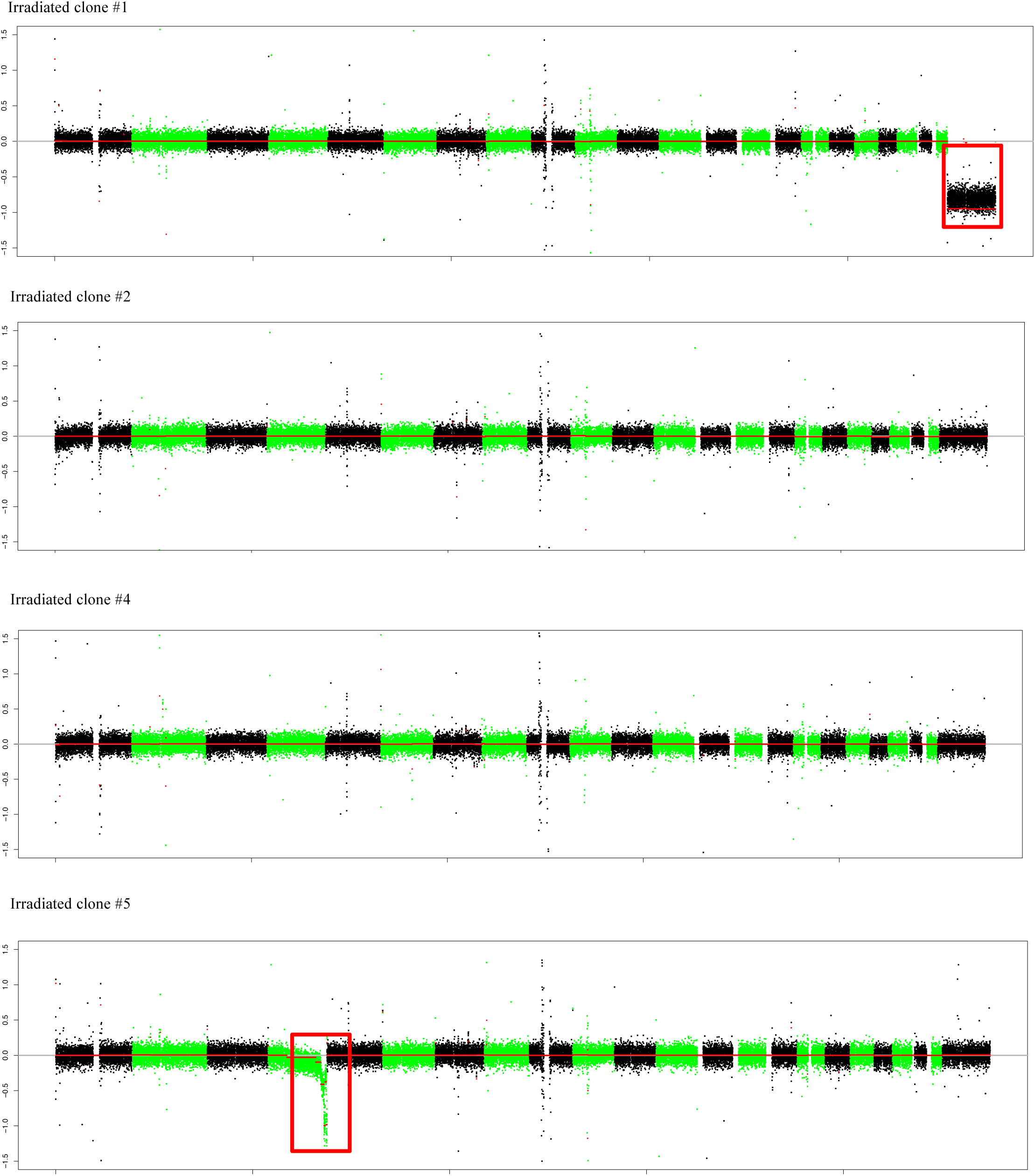
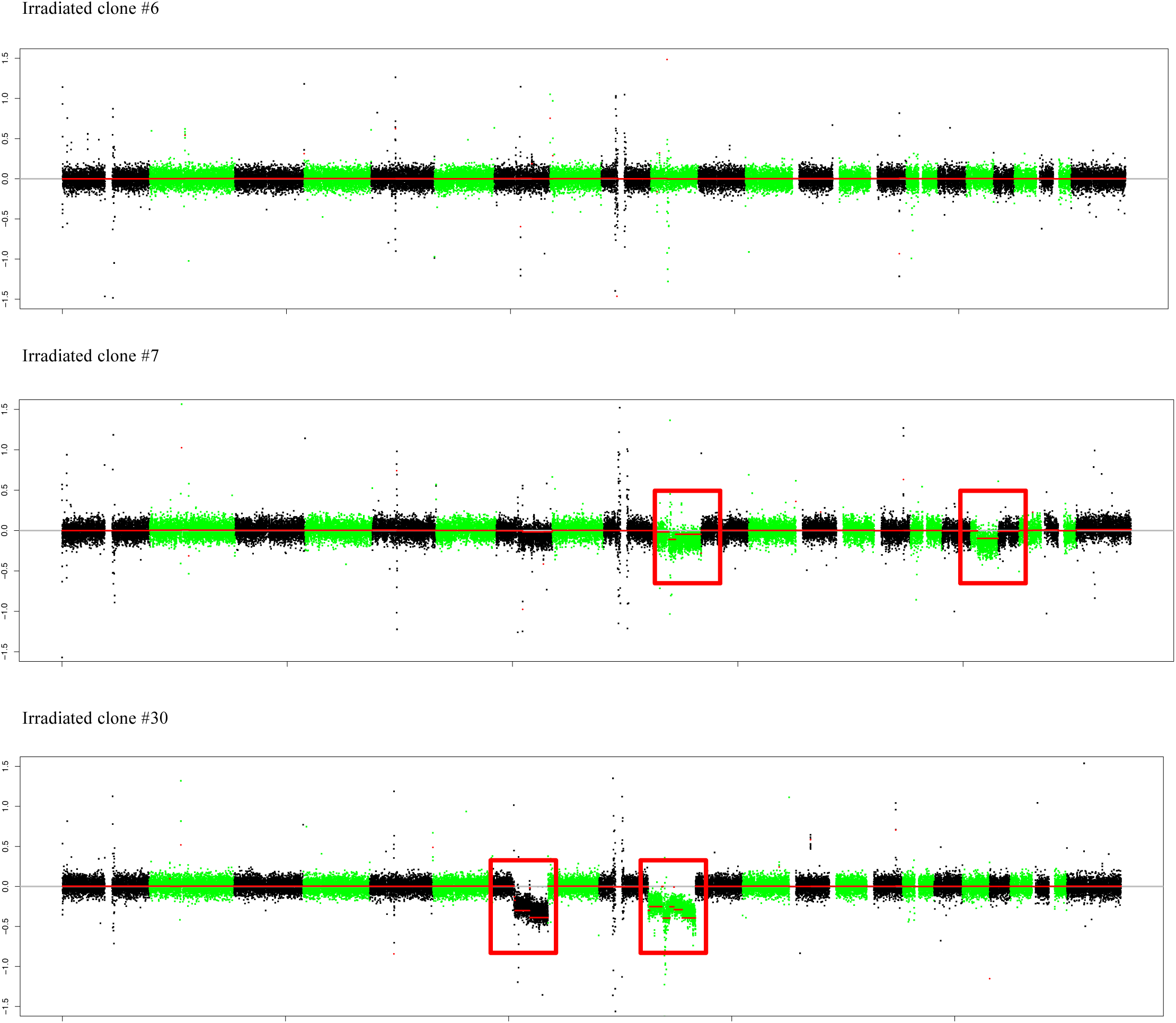
Copy-number plots for 11 RPE1 *TP53*-KO clones for which whole-genome sequencing was performed. Chromosomes with copy-number variations are highlighted.

**Supplementary Figure 6.**
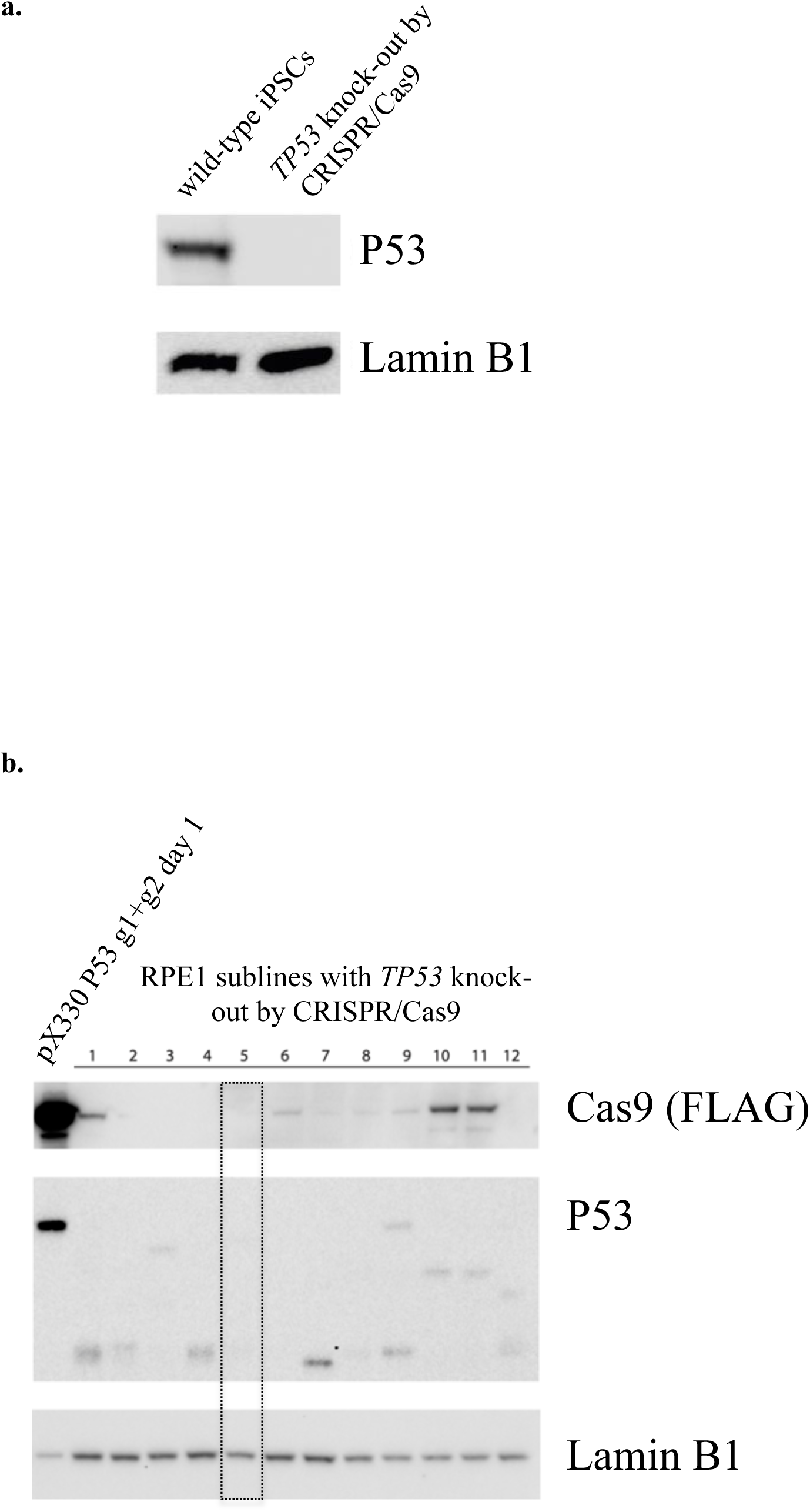
Western blot analysis of TP53 protein expression in wild-type induced pluripotent stem cells and after CRISPR-Cas9 mediated knock-out of *TP53* **(a)** and in wild-type RPE1 cells and sublines after *TP53* knock-out using CRISPR/Cas9 **(b)**. For all RPE1 *TP53* knock-out irradiation experiments, sub-line number 5 was used.

